# FliL has a conserved function in diverse microbes to negatively modulate motor output via its N-terminal region

**DOI:** 10.1101/2022.05.17.492383

**Authors:** Xiaolin Liu, Anna Roujeinikova, Karen M. Ottemann

**Affiliations:** Department of Microbiology and Environmental Toxicology, University of California, Santa Cruz, CA 95064; Infection and Immunity Program, Department of Microbiology and Department of Biochemistry and Molecular Biology, Monash Biomedicine Discovery Institute, Monash University, Clayton VIC 3800, Australia

**Keywords:** mechanosensor, motility, flagella, transmembrane

## Abstract

Bacterial surface sensing is often conferred by flagella. The flagellar motor protein FliL plays a key role in this process, but its exact role has been obscured by varying *fliL* mutant phenotypes. We reanalyzed results from studies on these *fliL* alleles and found they inadvertently compared mutants with differing length of the retained native N-terminal region, including the transmembrane helix (TM). We find that TM retention in the mutants that lack the native C-terminal domain results in loss of swimming and swarming motility, while alleles that completely lack the TM retain motility. We suggest FliL negatively regulates motor function via its N-terminal region, an observation that may relate to FliL function in mechanosensing.

## Introduction

Bacteria are skilled at sensing and adapting to their environments. One key environmental signal that they sense is surface proximity, which triggers a transition from the planktonic (single-cell, untethered) state to the surface-attached (sessile) biofilm state. The sessile-state cells are particularly important as they tolerate antimicrobial agents and cause numerous therapeutic and environmental problems (1), the key to solving which may lie in gaining control over the switch between the two states (2). Our current understanding is that planktonic bacteria sense surfaces to trigger expression of the genes needed to commence sessile growth, but the exact mechanism of this sensing is not clear. It has become appreciated that bacteria can employ their flagella as a direct surface sensor (2). The contact-induced mechanical hindrance to flagellar rotation has been suggested to lead to changes in torque generation, flagellar motor composition, and downstream physiological adaptations through a process called mechanosensing (3). However, the molecular mechanosignaling pathway that links the flagellum-surface contact to a cellular response is not yet known.

The flagellum is a complex nanomachine that is formed by over 30 proteins and has an ion motive force-powered motor at its base in the cell wall and cytoplasmic membrane (4). An influx of ions through the cytoplasmic membrane-embedded stator complexes (MotA, MotB) generates torque (5) that is applied to the cytoplasmic ring (FliM, FliN, FliG) of the rotor, making the rotor and the extracellular filament, attached to it via the hook, spin. Currently, it is thought that bacteria swimming in liquid have unimpeded flagella rotating without obstruction. When the cells begin to adhere to a surface, the flagellum-surface contact significantly increases the mechanical load on the flagellum, stalling the motor and triggering mechanosensitive signaling (3).

Studies have identified some flagellar proteins involved in the process of mechanosensing (6). One of these is FliL, a single-span membrane protein with a cytoplasmic N-terminus and a larger extracytoplasmic C-terminal region, the fold of which resembles an SPFH (stomatin/prohibitin/flotillin/HflK/C) domain found in proteins involved in the regulation of ion channel activities (7, 8). FliL associated with MotA and MotB, and weakly with FliG, in two-hybrid and pulldown assays (9). Recently, cryogenic electron tomography revealed that the extracytoplasmic domain of FliL forms a circle of rings, each coaxially sandwiched between MotA and the peptidoglycan-binding domain of MotB of a respective stator unit (7, 10), supporting the idea that FliL interacts intimately with the flagellar motor. The physiological function of FliL, however, remains mysterious. FliL has been suggested to be required for flagellar rotation, motor integrity, and/or surface sensing. The literature, however, is confusing because mutations in the *fliL* gene were reported to have different (and sometimes opposite) effects in different microbes. In some species, Δ*fliL* mutants were non-motile (7, 11-12), while in others they had enhanced or slightly decreased swimming motility (13, 14). *fliL* mutants have also been evaluated for swarming motility, a type of motility used in a more viscous environment and associated with surface sensing (15). As with swimming, Δ*fliL* mutants in some species showed severe swarming defects (16), while others showed enhanced swarming ability (14). Overall, these studies showed FliL is important for swimming and swarming motility, but it has been difficult to assess its exact mechanistic role and whether it plays similar roles in diverse microbes due to the variable phenotypes. Here we provide insight into this issue by identifying that previous work utilized different types of *fliL* mutants. We identify that a key variation is inclusion or exclusion of the cytoplasmic and transmembrane regions, and suggest the FliL N-terminal region acts as a motility inhibitor when retained without the extracytoplasmic C-terminal region.

## Results

*fliL* mutant phenotypes are reported to vary even within the same bacterial species (13, 14, 16, 17). We hypothesized that these divergent phenotypes occur due to differences in how the *fliL* gene was mutated, and what FliL parts were retained. We thus examined and categorized the previously-used mutations. We focused our study on bacterial species with one polar or peritrichous system, and excluded bacteria that have two types of flagellar systems (*Vibrio alginolyticus* and *Bradyrhizobium diazoefficiens* (8, 18)), because they each encode a FliL, but relations between these systems and their components are not perfectly clear. Mechanosensing is often studied using bacterial swarming behavior, which is observed using a narrow range of agar concentrations above that used for swimming (0.3%) but not exceeding 1% (15). We therefore selected strains with swarming ability information to be able to assess mechanosensing phenotypes. Our data set, compiled according to these criteria, contained *fliL* mutants from a total of six species belonging to Alphaproteobacteria (*Caulobacter crescentus* (17)), Betaproteobacteria (*Herminiimonas arsenicoxydans* (12)), Gammaproteobacteria (*Escherichia coli* (13, 14, 16), *Salmonella typhimurium* (13, 16), *Proteus mirabilis* (14)) and Campylobacterota (*Helicobacter pylori* (7)).

Analysis of the primary and secondary structure showed that FliL proteins from these bacteria share a common fold comprising a short, 12-28-residue cytoplasmic region at the N-terminus, a ∼23-residue transmembrane region (TM), and a ∼200-residue C-terminal extracytoplasmic domain that is connected to the TM by a variable-length linker (Fig. 1). The conserved structure suggests that these FliL proteins perform similar functions.

**Figure 1.**
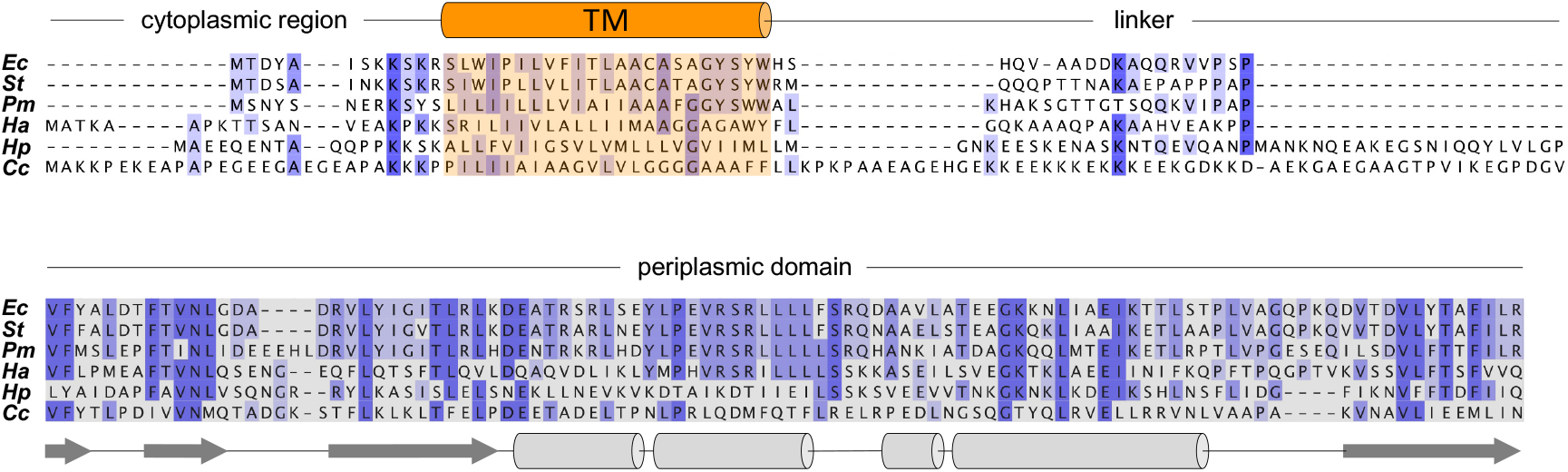
FliL proteins from different bacteria share a common general structure. Secondary structure-guided sequence alignment of FliL proteins from *Escherichia coli* (*Ec*), *Salmonella typhimurium* (*St*), *Proteus mirabilis* (*Pm*), *Herminiimonas arsenicoxydans* (*Ha*), *Helicobacter pylori* (*Hp*) and *Caulobacter crescentus* (*Cc*) is shown. Conserved amino acids are highlighted in blue, with darker color indicating higher degree of conservation. The predicted transmembrane (TM) helix is shown in orange, the periplasmic domain is shaded in grey. The secondary structure shown under the sequences was derived

We then analyzed the details of the Δ*fliL* mutants in different publications. When mutants are constructed, some regions at the 5’ or 3’ end of open reading frames are often retained by design or accident. We found that there was significant variation in the length of the N-terminal and C-terminal regions retained in the Δ*fliL* mutants. At the N-terminus, the variation can be divided into three types: loss of the TM (hereafter referred to as Δ*fliL1*); retention of the cytoplasmic region plus part of the TM (Δ*fliL2*); or retention of the cytoplasmic region plus the whole TM (Δ*fliL3*) (Fig. 2). At the C-terminus, the variation can be classified as complete loss, retention of about half of the FliL domain, or retention of a significant portion of the C-terminal domain.

**Figure 2.**
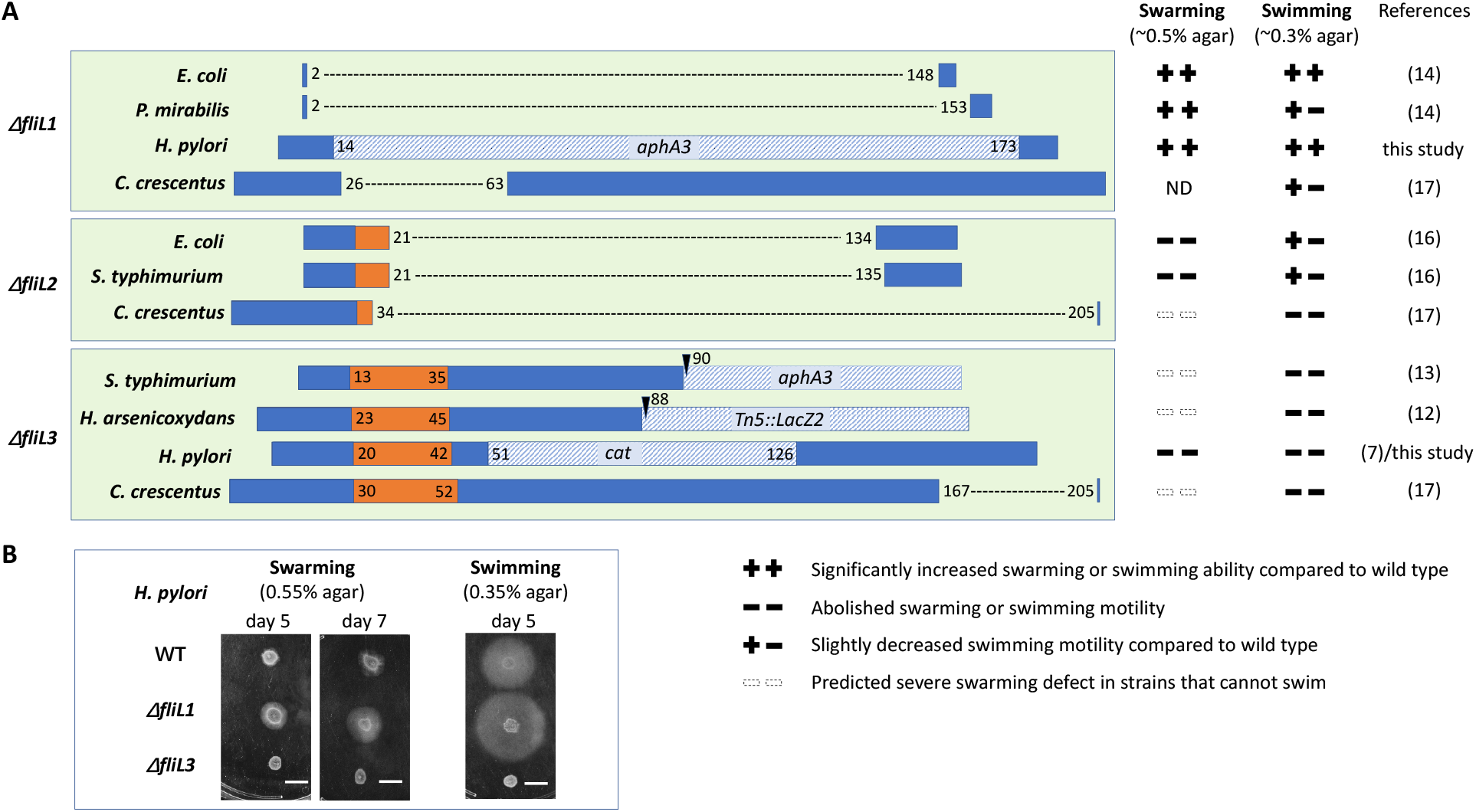
Comparison of *fliL* alleles with swarming and swimming phenotypes. (A) The diagram of regions deleted in different *fliL* alleles and their cognate phenotypes on swarming (∼0.5% agar) or swimming (∼0.3%) agar. The deleted regions are indicated with a dashed line, and any inserted antibiotic resistance genes are shown as regions with an oblique lines pattern plus the name of the gene. The transmembrane region of FliL is colored orange. The amino acid residue numbers at the region boundaries are shown. Δ*fliL1* alleles lack the entire TM region; Δ*fliL2* alleles lack part of the TM regions; Δ*fliL3* alleles retain the TM region. ND: Not Determined. (B) The swarming and swimming behavior of *H. pylori* wild type (WT), and Δ*fliL1* and Δ*fliL3* mutants after culturing for 5-7 days on 0.55% and 0.35% agar plates, respectively. The scale bar represents 1.0 cm.

We next compared these various *fliL* allele types with respect to their phenotypes for swarming or swimming motility. When we mapped the Δ*fliL* mutants to their phenotypes on swarming motility, we observed variation that correlated with the type of N-terminal mutation. Specifically, alleles that retained all or part of the TM (Δ*fliL2* and Δ*fliL3*) showed severe swarming defects (Fig. 2). In contrast, deletion of the entire TM (Δ*fliL1*) caused no swarming defects, and even resulted in swarming colonies that were larger than those of wild type. There was no obvious correlation with types of C-terminal mutations. These results suggest that the function of FliL in surface sensing involves its N-terminal region, including the TM, such that alleles that retain even a portion of the TM inhibit swarming motility.

We next analyzed variation in swimming behavior. As with swarming, Δ*fliL* mutant alleles that retained the full TM (Δ*fliL3*) were non-motile (Fig. 2), alleles with no TM (Δ*fliL1*) generally retained swimming motility, while alleles with a partial TM (Δ*fliL2*) typically showed an intermediate swimming phenotype. These results further suggest that FliL controls motility via its TM region. The intermediate phenotypes of Δ*fliL2* may be the consequence of a reduced efficiency of membrane insertion and hence lower occupancy of FliL at the motor due to the shorter hydrophobic membrane-targeting sequence.

The Δ*fliL* mutant previously constructed in *H. pylori* was non-motile and a *fliL3* allele (7) (Fig. 2A). We experimentally tested our hypothesis that the role of FliL in surface sensing and motility is associated with its TM by constructing an *H. pylori* Δ*fliL1* mutant lacking the entire TM (Fig. 2A). Consistent with our expectations, and in line with the previous report on *E. coli* Δ*fliL1*, the Δ*fliL1* mutant in *H. pylori* showed even stronger motility than wild type on both swarming and swimming soft agar plates (Fig. 2B).

## Summary

Our analysis suggests that FliL negatively regulates both swimming and swarming motility via its N-terminal region, including the TM helix. It’s not yet clear how it exerts negative control on the motor. A clue may come from the observation that motility was restored in the Δ*fliL2* mutants retaining the N-terminal region by extragenic suppressor mutations in the plug region of *motB* (9, 11). Given that FliL and MotB are close to each other in the motor and interact in vitro (7, 9, 10), we propose that the N-terminal FliL region interacts with MotB in the Δ*fliL2* mutants so as to prevent plug opening, block ion flow and inhibit both swarming and swimming motility; in wild-type FliL, the C-terminal domain may act in mechanosensing and regulate the N-terminal region. This new appreciation for the N-terminal region of FliL provides a framework to understand how the flagella may transduce mechanical signals. Furthermore, our analysis of the Δ *fliL1* allele appears to put an end to the long-standing argument about the requirement of this gene for motility by showing that FliL is not an essential component of the flagellar motor. This explains why this gene avoided detection by classical loss-of-function genetic analysis for a long time.

## Data Availability

The information about regions deleted in the *fliL* mutants and cognate phenotypes on swarming and swimming was collected from published articles as cited. The sequence alignment was performed using PRALINE (https://www.ibi.vu.nl/programs/pralinewww/) and edited in Jalview. The secondary structure and transmembrane regions were predicted using YASPIN and TMHMM (http://www.cbs.dtu.dk/services/TMHMM/), respectively. The Δ*fliL1* in *H. pylori* SS1 background was constructed using natural transformation. The first 40 bp of *fliL* and 500 bp upstream from its start codon was amplified using primers P212 and P213 (P212: cgggatccagagagtcgttttttaacaccttg, P213: acatagtatcgttgggggttgttgcgc).

The last 32 bp from stop codon and 500 bp downstream of *fliL* was amplified using primers P216 and P217 (P216: gatgaattgttttagatgtctttttcactgattttatcatcc, P217: acgcgtcgacgaagtttgattttatttatttagaatttgttatcaaaa). An *aphA3* cassette that confers kanamycin resistance was amplified by primer pairs P214/P215 (P214: caacccccaacgatactatgttatacgccaact, P215: cagtgaaaaagacatctaaaacaattcatccagtaaaatataatattttattttct). The three amplicons were fused by overlap extension PCR using primer pairs P212/P217 and the resulting product was used to transform *H. pylori*. The positive colony screened by kanamycin was confirmed by sequencing. Overnight cultures of *H. pylori* were prepared in Brucella Broth (BB) plus 10% (v/v) heat-inactivated fetal bovine serum (FBS). Swimming and swarming behavior of *H. pylori* were tested by adjusting the overnight culture to OD_600_ of 0.15 and then incubating in BB plus 2.5% (v/v) FBS containing 0.35% and 0.55% Bacto agar, respectively, for 5-7 days.

## ACKNOWLEDGMENTS

The described project was supported by the National Institute of Allergy and Infectious Disease (NIAID) grant 1R01AI164682-01 to K.M.O, the Australian Research Council grant DP210103056 to A.R., and a student fellowship (No. 201904910692) from the China Scholarship Council to X.L. The funders had no role in study design, data collection, and interpretation, or decision to submit the work for publication.

